# *Mycobacterium tuberculosis* grows linearly at the single-cell level with larger variability than model organisms

**DOI:** 10.1101/2023.05.17.541183

**Authors:** Eun Seon Chung, Prathitha Kar, Maliwan Kamkaew, Ariel Amir, Bree B. Aldridge

## Abstract

The ability of bacterial pathogens to regulate growth is crucial to control homeostasis, virulence, and drug response. Yet, we do not understand the growth and cell cycle behaviors of *Mycobacterium tuberculosis* (Mtb), a slow-growing pathogen, at the single-cell level. Here, we use time-lapse imaging and mathematical modeling to characterize these fundamental properties of Mtb. Whereas most organisms grow exponentially at the single-cell level, we find that Mtb exhibits a unique linear growth mode. Mtb growth characteristics are highly variable from cell-to-cell, notably in their growth speeds, cell cycle timing, and cell sizes. Together, our study demonstrates that growth behavior of Mtb diverges from what we have learned from model bacteria. Instead, Mtb generates a heterogeneous population while growing slowly and linearly. Our study provides a new level of detail into how Mtb grows and creates heterogeneity, and motivates more studies of growth behaviors in bacterial pathogens.

## Introduction

Tuberculosis (TB) remains challenging to treat, requiring lengthy multidrug treatment due to heterogeneity in host response and the behaviors of individual *Mycobacterium tuberculosis* (Mtb) bacilli^1–6^. Drug-tolerant subpopulations of Mtb exhibit different characteristics in growth, metabolic state, and gene regulation^7–13^. Therefore, the ability of Mtb to develop heterogeneity in many cellular processes, including growth behaviors, is thought to be a major obstacle in developing shortened therapies^3, 14–16^. Phenotypic switching to a slow growth state to create persistence has been observed in other bacteria, including *Escherichia coli* (*E. coli*)^17^ and Salmonella^18, 19^. Despite the critical role of growth variation in the ability of bacteria to tolerate drug treatment, the basic growth behaviors and heterogeneity in growth characteristics of Mtb are unknown, making it challenging to study how growth variation is created and maintained in this pathogen.

Single-cell level growth studies of mycobacteria have focused on the non-pathogenic species *Mycobacterium smegmatis* (*M. smegmatis*), which is faster growing and larger than Mtb^20–25^. Mycobacteria elongate from their poles, and *M. smegmatis* grow and divide asymmetrically, creating variation in growth behaviors between sister cells. *M. smegmatis* elongate more from their old poles, and the sister inheriting the old pole (accelerator cell) is longer at birth and grows faster than its sister (alternator cell). Our understanding of growth behaviors in *M. smegmatis* is based on data from live-cell and fixed-cell imaging, and practical challenges have made it difficult to perform analogous live-cell imaging experiments in Mtb due to its slow-growing nature, small size, and the requirement for handling Mtb in a biosafety level-3 conditions. As a result, our understanding of growth behaviors in Mtb is based mainly on *M. smegmatis* and fixed-cell imaging in Mtb^20, 21, 23, 25–28^. There is evidence that Mtb can grow and divide asymmetrically, but it is not clear that the asymmetry is as consistent as it is in *M. smegmatis*^29, 30^. It may be that the growth behaviors of Mtb are not very similar to those of *M. smegmatis,* given the difference in time scales and dimensions. The doubling time of *M. smegmatis* ranges from 3 hours (fast-growing condition) to 5 hours (carbon-limited conditions)^24, 31–33^. In contrast, Mtb doubles every ∼18h in rich medium, and the doubling times increase in host-mimicking conditions^3, 34–36^. Reflecting these dramatic differences in growth dynamics, the cytoplasmic ribosome density of *M. smegmatis* is significantly higher than that of Mtb^30, 37^. The fundamental differences in size, growth rate, and ribosome density of *M. smegmatis* and Mtb suggest that understanding the growth behaviors of Mtb requires a direct study of Mtb using time-lapse imaging rather than a transfer of learning from *M. smegmatis*.

Though cells double in number in each generation with their population growing exponentially, it is unclear a priori how a single cell grows. Determining how cells grow is crucial as the growth mode constraints which molecular mechanisms may be involved in cell growth^38–40^. Advances in microfluidic devices, microscopy, and image analysis have launched a new era of single-cell measurements at high-throughput levels, leading to developments in single-cell analysis methods and quantitative models of cell growth behaviors ^41–50^. These single-cell data point to most bacteria growing exponentially^41, 42, 51–56^. Even archaea, which belong to a different domain of life, were reported to grow exponentially^57^. The exponential growth of single cells is explained by the self-producing ability of ribosomes and the fact that ribosomes produce other proteins^39^. Recent analysis methods have shown *E. coli* to grow super exponentially in length during the later stages of the cell cycle^58–60^. A biphasic growth mode was identified recently in *Bacillus subtilis* (*B. subtilis)*, where cells grow linearly until they reach a certain size (sizer) after which they grow exponentially for a fixed time (timer) until they divide^61^. *C. glutamicum*, a tip grower like mycobacteria, has been observed to undergo asymptotic linear growth, in which the rate-limiting step for growth is the synthesis of the polar cell wall^62^. Complex growth rate trends are observed in eukaryotic organisms such as the fission yeast *Schizosaccharomyces pombe*, where growth rates are modulated to maintain proteome concentration homeostasis^40^. Thus, the study of the mode of growth becomes crucial for understanding homeostasis and cell cycle regulation mechanisms.

In this study, we characterize the fundamental single-cell growth characteristics and growth mode of Mtb for the first time. Using time-lapse imaging, we measured single-cell growth and cell cycle parameters to describe Mtb growth mode and detailed growth behaviors, including cell size parameters and the origins of growth heterogeneity. We show that Mtb grows and divides asymmetrically and exhibits high levels of heterogeneity in cell size, interdivision time, and elongation speed. However, unlike *M. smegmatis*, accelerator and alternator subpopulations do not have different growth behaviors in Mtb. In unbuffered and acidic conditions, we discovered that Mtb exhibits a nearly linear growth mode at a single-cell level. Using pulse-label experiments of new cell wall synthesis, we find that in about half of the cells, elongation occurs from both poles from birth without a time lag. Together, our study provides a new quantitative framework to study growth behaviors and variation in Mtb.

## Results

### Live-cell imaging to measure Mtb growth behaviors

In the past few decades, advances in time-lapse imaging and microfluidics technologies have transformed our ability to characterize cellular growth behaviors at the single-cell level^41, 42, 47, 48, 52, 63^. However, the single-cell behaviors of Mtb have remained elusive because of the practical challenges of using these technologies on this slow-growing pathogen. The highest throughput microfluidics for studying cell growth in rod-shaped bacteria require that the bacteria are loaded into thin channels^52, 64, 65^. Mycobacteria cannot be loaded into these channels because they are coated in a thick mycolic acid layer in their cell wall, making them too sticky for these devices (Extended Data Fig. 1a). Instead, we load them into shallow “rooms”. This ensures a consistent growth medium (via diffusion from a constant flow channel that interfaces with the rooms) and helps maintain focus for live-cell imaging (Supplementary File1). Similar microfluidic chambers are used to study *Halobacterium salinarum* (*H. salinarum*)^57^, a model archaeon that is sensitive to mechanical stresses, and *Corynebacterium glutamicum* (*C. glutamicum*), a tip-grower like mycobacteria^62^. There are other designs of microfluidic devices that are developed for mycobacteria, including using geometries that creates gradient patterns of drugs^66^. In simple microfluidic devices with a media flow channel and attached shallow rooms, mycobacteria have the freedom of movement necessary for polar growth and to divide in their v-snapping (forming a V-shape during the process of bacterial cell division) pattern (Extended Data Fig. 1b,c)^67, 68^. The growth pattern and stickiness of the cells create cell clumps that make many cells unsuitable for annotation (Extended Data Fig. 1a). Mtb bacilli (2-4 µm in length, 0.34 ± 0.03 µm in diameter) are smaller than *M. smegmatis* (3-5 µm, 0.58 ± 0.05 µm in diameter), making it harder to maintain cells in focus throughout the growth cycle and clumps can easily form in the channels, which are 1-1.2 um in depth^30, 33, 37, 69, 70^. Mtb grows very slowly (doubling time of 16∼18 hours during imaging in rich growth conditions, compared to other bacterial species such as *M. smegmatis* with 3 hours and *E. coli* with 20 minutes of doubling time, respectively) and must be handled in a biosafety level-3 laboratory, making time-lapse imaging experiments resource intensive. Furthermore, the irregular and small shape of Mtb bacilli poses difficulties for automated segmentation and tracking software, making manual annotation a necessary task (Extended Data Fig. 1b). Collectively, these obstacles have made it more challenging to obtain live-cell imaging data of Mtb.

**Fig. 1.**
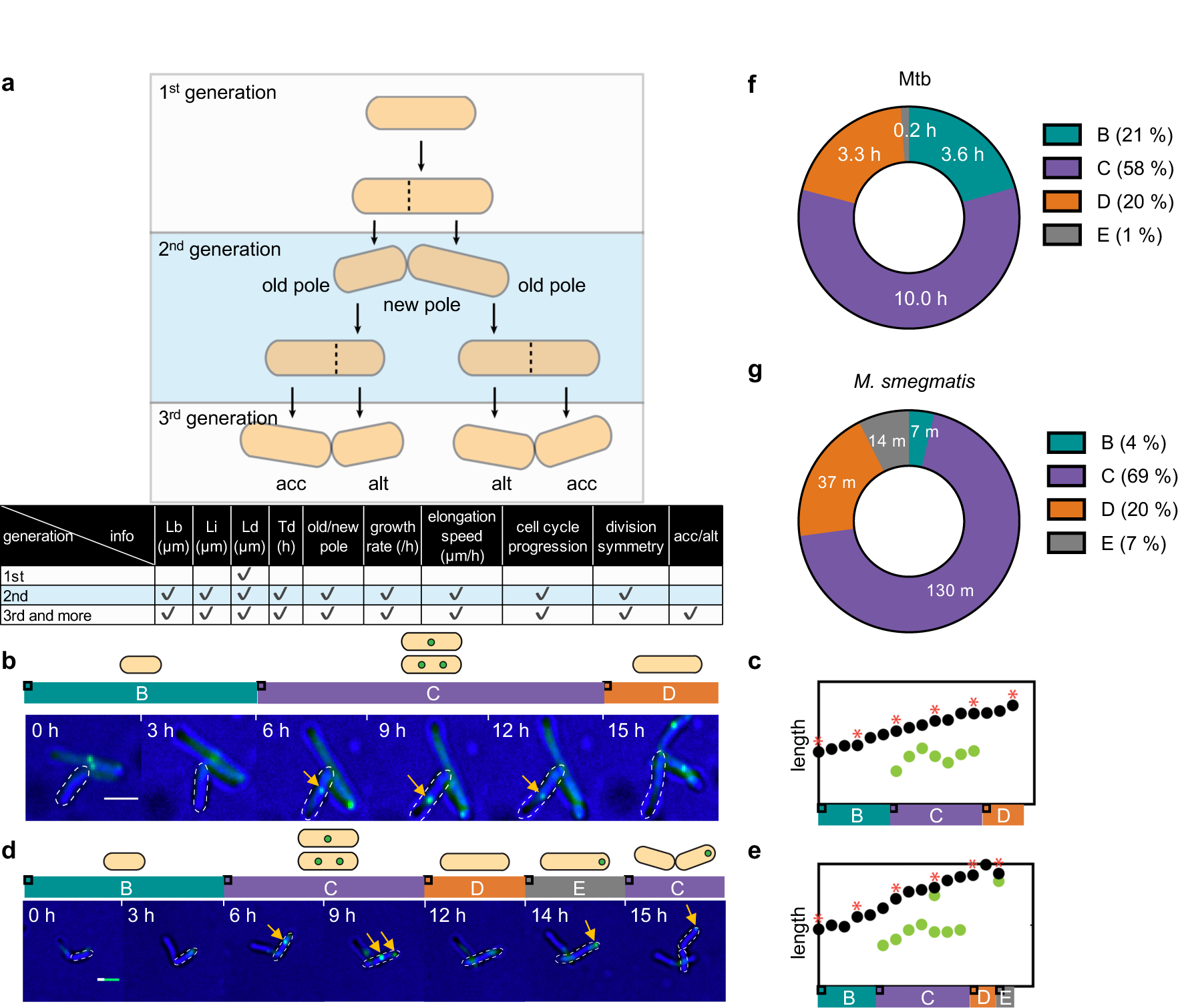
Measurement of growth characteristics in Mtb (movie set A) **a**, Schematic of growth parameters derived from Mtb at different generations by time-lapse imaging. Cells loaded into the device (1st generation) are used to establish the pole age for the next generation. However, they are otherwise excluded from analysis because the entire cell cycle is not observed. In the 2nd and subsequent generations, all growth features can be determined, including length at birth, initiation of DNA replication, and division (Lb, Li, and Ld), interdivision time (Td), growth rate (*λ*), elongation speed 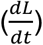, and identification of the old pole and the new pole. In the 3rd (and subsequent) generations, we are also able to determine whether a cell is an accelerator (acc) (n = 173) or an alternator (alt) (n = 130). In the baseline experiments (movie set A), we determined cell cycle timing at the single-cell level using an SSB-GFP reporter strain of Mtb. The total annotated cell number is 363 single cells. **b**, Determination of cell cycle state using SSB-GFP. Single-stranded binding protein (SSB) binds to replication forks^79, 80^, so the presence of green foci using an Mtb reporter strain with SSB-GFP indicates that chromosome replication is in progress. There are no visible foci before and after replication (B and D periods, respectively). One or two foci appear during replication (C period). Yellow arrows point to foci. Scale bar = 2 µm. **c,** Single-cell traces of SSB-GFP localization with hourly time points of the cell shown in (**b**). The distance between the x-axis and the black circle indicates cell length. Green circles indicate SSB-GFP foci localization along the cell body. Asterisks (*) correspond to the images shown in (**b**). **d**, Annotation of cell cycle state in a cell with an E period. A small population of cells (11 %) exhibits a new round of replication that starts before cell division (foci-positive), termed the E period. Daughter cells inheriting the foci enter directly into the C period. Yellow arrows point to foci. Scale bar = 2 µm. **e**, Single-cell traces of SSB-GFP localization with hourly time points of the cell shown in (**d**). The distance between the x-axis and the black circle indicates cell length. Green circles indicate SSB-GFP foci. Asterisks (*) correspond to the snapshots shown in (**d**). **f**,**g**, Cell cycle timing in Mtb (**f**) and *M. smegmatis* (**g**). The average time and proportion of each cell cycle period (B: pre-replication, C: DNA replication, D: post-replication, E: new DNA replication after a D period but before division). The *M. smegmatis* data are from a previous study^24^.

We overcome these challenges by optimizing protocols and using a custom microfluidic device to achieve long-term time-lapse imaging of Mtb in a biosafety-level 3 laboratory. We used devices previously designed to study *M. smegmatis* to ensure freedom of movement in polar growth and v-snapping^22, 24^. We observed that, whereas *M. smegmatis* grows with a new medium constantly flowing in the microfluidic devices, Mtb enters growth arrest. Culture filtrate is required to enable some subpopulations of Mtb to grow^71–73^, so we supplement the new medium flowing into the device with culture filtrate at a ratio of 1:1 to avoid growth arrest. With these protocols, we were able to achieve a consistent growth pattern in Mtb over four days of imaging with a doubling time (∼17h) that is consistent with the doubling times of Mtb in bulk culture^74–76^. We used the simple imaging platform to record Mtb cultures growing for four days with images captured every hour (see Methods, movie set A). Annotation of these images allowed us to calculate many growth features at the single-cell level, including cell length at birth (Lb) and division (Ld), growth rate, and elongation speed (Fig. 1a). By tracking cell lineage, we identified sister cells and annotated which sister inherited the old and new poles (accelerator and alternator cells, respectively) (Fig. 1a). The old pole is the pole that daughter cells inherit from the mother cell, while the new pole is the pole that previously formed the septum in the mother cell and transitions into a new pole. To determine cell cycle timing, our base movie set was made with the Mtb strain CDC1551 carrying a fluorescent reporter of active DNA replication via a GFP-tag on an episomal copy of single-stranded binding protein (SSB) (Fig. 1b,d)^77^. Cells undergoing active DNA replication (C period) have visible GFP foci because SSB binds to single-stranded DNA at replication forks. Cells that are in the pre- or post-replication phases (B and D periods, respectively) do not have a visible GFP focus (Fig. 1b,c and Extended Data Fig. 2a-c)^12, 24, 77^. A small population of Mtb cells (11% of total cells) enter another round of replication period before septation; this has also been observed in *M. smegmatis*, and we label this as the E period (Fig. 1d,e and Extended Data Fig. 2d,e)^24, 78^. By annotating when SSB-GFP foci appear and disappear, we can calculate the length of the B, C, D, and E periods for single cells and the average duration of each period in the total population (Fig. 1f,g). We performed time-lapse imaging of these cells in biological triplicate in ∼70 chambers per movie, resulting in ∼2700 cells captured in the movies, of which we were able to fully annotate 363.

**Fig. 2.**
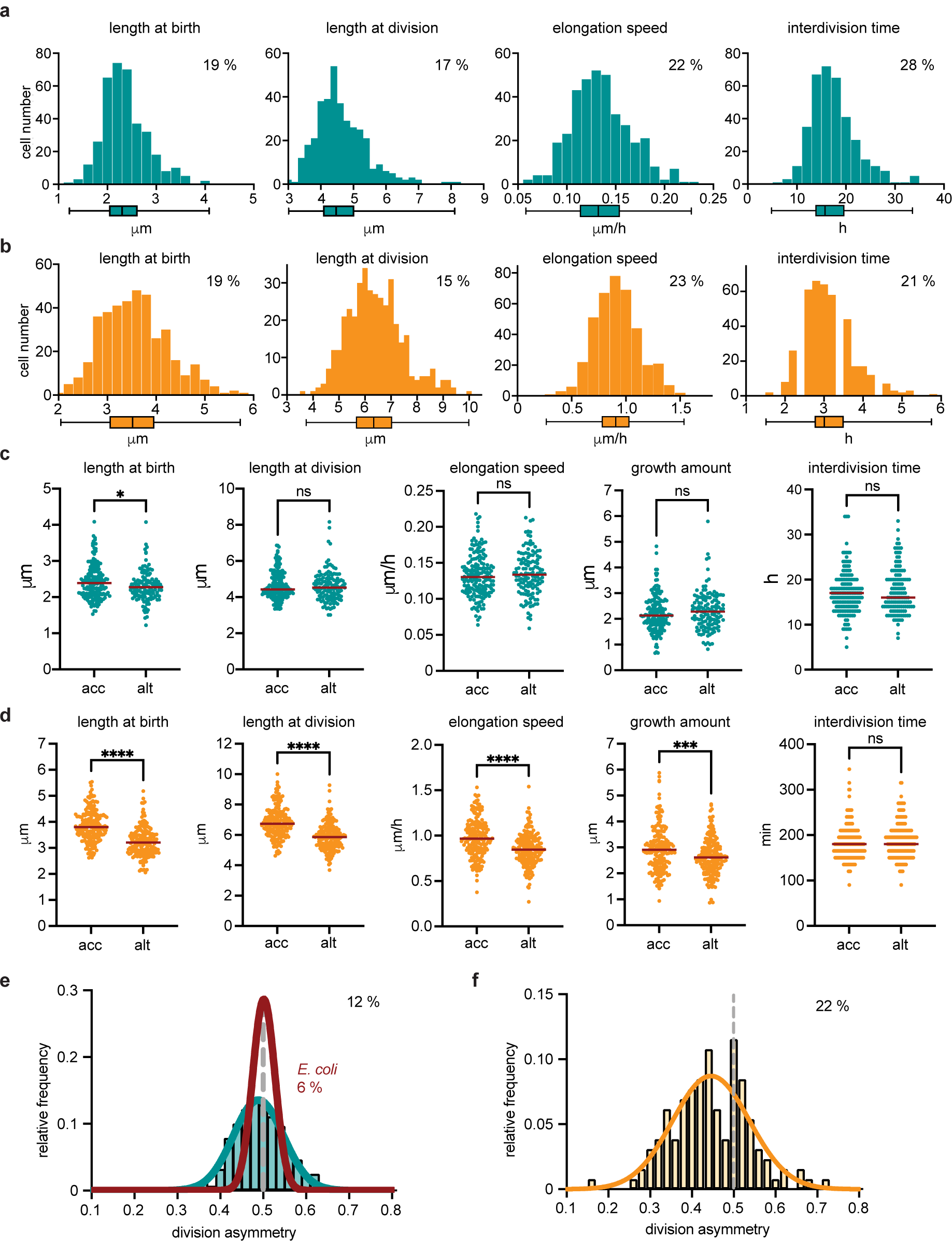
Growth properties in Mtb and *M. smegmatis*. **a,b**, Distributions of Lb and Ld, Td, and elongation speed 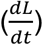 are shown in both (**a**) Mtb (n = 363; movie set A) (green) and *M. smegmatis* (n = 391) (orange) (**b**), respectively. The coefficient of variation is shown in the upper right corner of each plot. The *M. smegmatis* data are from a previous study^24^. **c**,**d**, Growth property distributions are compared between acc and alt cell in Mtb (n of acc = 173, n of alt = 130) (green) and *M. smegmatis* (n of acc = 193, n of alt = 188) (orange) (**d**). Lb and Ld, elongation speed, growth amount, and Td are compared between acc and alt, respectively. Horizontal lines mark the median value for each sample. The *M. smegmatis* data are from a previous study^24^. P-values were calculated with a Mann-Whitney test (p-value < 0.05, *; < 0.001, ***; < 0.0001, ****). Mann-Whitney *U* = 9542, 10908, 10592, 10419, 11050 in Mtb and 7576, 8326, 11712, 14335, and 16207 in *M. smegmatis*, respectively. **e**,**f**, Distributions of division asymmetry in Mtb, *M. smegmatis* and *E. coli*. The distributions of division asymmetry in Mtb (green) (**e**) and *M. smegmatis* (orange) (**f**) are calculated as (Lb of alt cell (daughter cell) / Ld of mother cell). The division symmetry of *E. coli* (red) is calculated as (Lb of daughter cell / Ld of mother cell) (**e**). The *M. smegmatis* and *E. coli* data are from previous studies^24, 57^. The dashed lines at 0.5 indicate perfect symmetry. The CVs of Mtb, *E. coli*, and *M. smegmatis* are 12 %, 6 %, and 22 %, respectively.

### Cell cycle timing in Mtb

Mtb’s doubling time (∼17 hours) is more than five times slower than it is in *M. smegmatis* (∼3 hours) (Fig. 1f,g), so we would not expect that cell cycle timing is similar in both species (either in absolute time or proportionally). We tracked when SSB-GFP foci appeared and disappeared (Fig. 1b-e and Extended Data Fig. 2) to determine the duration of each cell cycle period and their proportion of the total cell cycle (Fig. 1f). On average, Mtb spends 3.6 hours in the pre-replication period (B), 10 hours in the replication period (C), 3.3 hours in the post-replication period (D), and 0.2 hours of the period where a new replication round begins before division (E period, see Fig. 1d,e and Extended Data Fig. 2d,e). Together, the B-C-D-E periods comprise 21%, 58%, 20%, and 1% of total cell cycle duration, respectively (Fig. 1f). We found that the cell cycle period was not proportionally similar in Mtb and *M. smegmatis* (Fig. 1f,g). The C period is proportionally shorter in Mtb, and division occurs roughly in the middle of the non-replication period so that B and D are nearly equal. In contrast, in *M. smegmatis*, the B period is much shorter than the D period. A larger proportion of cells (47%) initiate a new round of replication before division so that their daughters are born in C and do not have a B period^24, 77, 78^.

### Mtb cell lengths, interdivision times, and elongation speeds are heterogeneous

Next, we evaluated the growth characteristics of Mtb. To ensure that the microenvironment of the microfluidic chambers and light exposures during imaging did not fundamentally alter growth behaviors compared to those measured by other studies, we compared doubling times and cell lengths to those reported in the literature. In general, the length of Mtb bacilli ranges from 2 to 4 µm *in vitro*^69^. The cell lengths from our movies were consistent with this reported range with median lengths at birth and division of 2.3 µm and 4.5 µm, respectively (Fig. 2a). The background strain of Mtb (CDC1551) doubles every 17.1 ± 4.2 hours^75^, which is similar to the 16 h we observed as the median interdivision time (doubling time) of the CDC1551 SSB-GFP reporter strain by time-lapse imaging (Fig. 2a). In addition, we evaluated growth characteristics of cells born at different times over the four-day movies and did not observe significant variation between cells born early or late in the movie or among replicates (Extended Data Fig. 3).

**Fig. 3.**
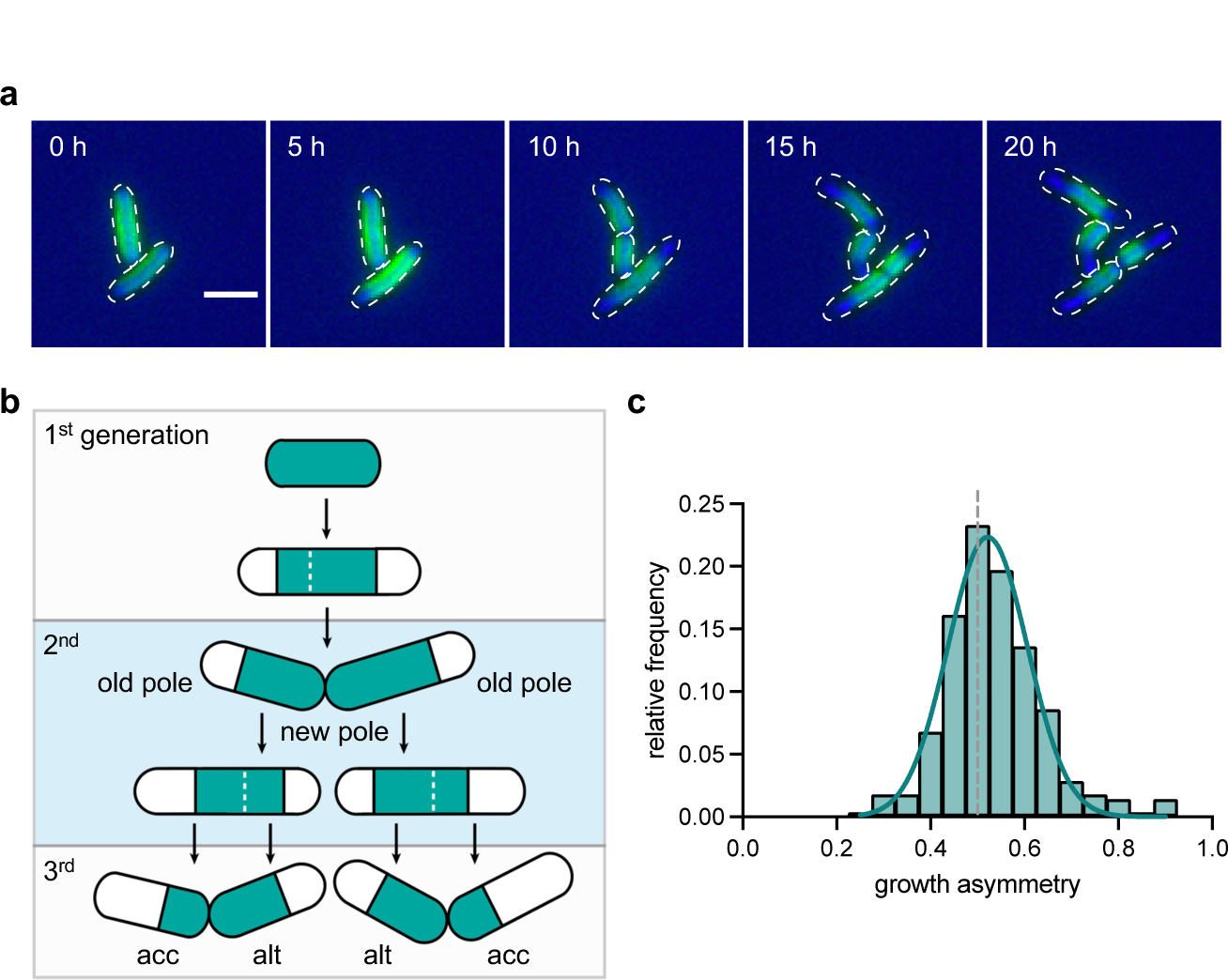
Polar growth and growth symmetry in Mtb. (measured in movie set B)**. a**, 5-hour-interval image sequence of FDAA-labeled Mtb cells. Individual cells were fully labeled with HADA at the beginning of the time-lapse imaging. The unlabeled area at the poles indicates a newly grown cell wall since the start of the imaging. Scale bar, 2 µm. **b**, Schematic diagram of polar growth quantification using FDAA-labeling Mtb. The blue area within the cell indicates FDAA (HADA) labeling. The white (unlabeled) area at the poles indicates the new cell wall. The white dashed line indicates the septum formation. **c**, Distribution of growth symmetry in Mtb. The histogram is fit with a Gaussian. The gray dashed line (score of 0.5) represents symmetric growth. (mean value = 0.58, n = 147).

Studies of mycobacterial growth characteristics have been performed primarily in *M. smegmatis*, a model organism for mycobacterial cell biology^24, 25, 33, 81, 82^. In *M. smegmatis*, growth and division are asymmetric, giving rise to more heterogeneity in growth characteristics than in other bacterial species^4, 24–27, 83^. To describe Mtb growth characteristics and compare levels of heterogeneity to *M. smegmatis*, we compared distributions of Lb and Ld, Td, and elongation speed in Mtb to *M. smegmatis* (Fig. 2a,b). We observed that Mtb growth is heterogeneous to the same degree as in *M. smegmatis* for birth length (Mtb 19 % and *M. smegmatis* 19 % in CV, p-value = 1) and elongation speed (Mtb 22 % and *M. smegmatis* 23 % in CV, p-value = 0.41). Heterogeneity is increased in Mtb compared to *M. smegmatis* for division length (Mtb 17 % and *M. smegmatis* 15 % in CV, p-value = 0.018) and interdivision time (Mtb 28 % and *M. smegmatis* 21 % in CV, p-value = 1.42 x 10^-7^).

### Accelerator and alternator Mtb cells exhibit similar growth properties

Growth and division asymmetry in *M. smegmatis* leads to a deterministic heterogeneity in growth properties. For example, the accelerator cells are longer, elongate more, and elongate faster than the alternator cells (Fig. 2d)^24–27^. Because Mtb exhibits slightly more heterogeneous growth behaviors in cell sizes and interdivision timing than *M. smegmatis*, we speculated that differences in growth characteristics between accelerator and alternator cells would be amplified in Mtb. Instead, we found no significant difference in their behaviors except for a slight difference in cell size where accelerator cells are born longer than alternator cells (Fig. 2c, p-value, Lb 0.02, Ld 0.66, elongation speed 0.39, growth amount 0.27, interdivision time 0.80).

### Mtb division asymmetry is variable

Because accelerator and alternator cells exhibit similar growth behaviors in Mtb, including cell size, we speculated that division might be relatively symmetric compared to *M. smegmatis*. We quantified the division asymmetry level by calculating the ratio of the length of the alternator cell at birth to the sum of the length of both sisters at birth. Accordingly, the distribution of division asymmetry would center around 0.5 for symmetric division and below 0.5 when the alternator cells are smaller than accelerator cells (Fig. 2e,f). Reflecting the relatively minor difference in birth sizes between accelerator and alternator cells in Mtb compared to *M. smegmatis*, we observed that the Mtb distribution is centered near 0.5 (0.49), whereas the *M. smegmatis* distribution is centered around 0.45 (Fig. 2e,f). We noted that the distribution of division symmetry for Mtb, though narrower than in *M. smegmatis* (CV of 12% vs. 22%, p-value = 1 × 10^−17^, Fig. 2f), is significantly more variable than in *E. coli* (CV 6%, p-value = 5 × 10^−29^) (Fig. 2e). Together, these data suggest that growth heterogeneity is less polar in Mtb than it is in *M. smegmatis*, but that division is not as symmetric as in model rod-shaped bacteria like *E. coli*.

### Mtb growth is slightly asymmetric with more elongation from the old pole

We next investigated whether growth was asymmetric in Mtb. Mycobacterial elongation is polar, and growth is faster (and earlier) from the old pole compared to the new pole in *M. smegmatis*^25, 26, 29, 84–86^. In Mtb, fixed-cell imaging of growth patterns using pulse-chase experiments of a labeled cell wall with amine-reactive dyes or fluorescent D-amino acids (FDAAs) has suggested that Mtb growth is also asymmetric but less asymmetric than in *M. smegmatis*^26, 27, 29, 86, 87^. However, it has been challenging to quantify the asymmetric growth pattern and determine whether there is more growth from the old pole than the new pole without time-lapse imaging. By tracking cell surface features, Hannebelle and colleagues observed NETO (new end take-off), that is the old pole elongates before the new pole elongates, in both Mtb and *M. smegmatis*^86^. To measure growth symmetry throughout entire division cycles in Mtb, we followed Mtb growth by time-lapse imaging for ∼6 days (140 hours) after pulse labeling Mtb with a blue fluorescent d-alanine (HADA) for 24 hours (Fig. 3a,b, movie set B). Using the HADA label as a marker of old cell wall material, we then annotated growth from the new and the old poles in cells born during the movie (so that we could establish pole age) from birth to division (n = 248) (Fig. 3a,b). Unlabeled cell wall regions at the poles marked the growth at each (old and new) pole (Fig. 3a,b). The start of growth and its amount can often be hard to characterize because of cell clumping (Extended Data Fig. 1a), errors due to finite resolution limit during imaging, and diffusion of the HADA label to the newly formed region. Thus, we used mathematical models to quantify the amount of growth using the time course of growth annotations. Based on previous results of linear growth at the old pole and bilinear growth at the new pole^86^, we fit linear and bilinear models to characterize the length growth at each pole as a function of time from birth (see Methods and Extended Data Fig. 9). A linear model implies growth at constant elongation speed from the beginning of the cell cycle, whereas a bilinear model implies a lag period where the cell pole does not grow following which the cell grows at a constant speed (Extended Data Fig. 4). Only one of the two models is chosen as the mode of growth for each pole of an individual cell. The choice is made based on the Akaike information criterion (*AIC*) and the Bayesian information criterion (*BIC*) metrics (see Methods).

**Fig. 4.**
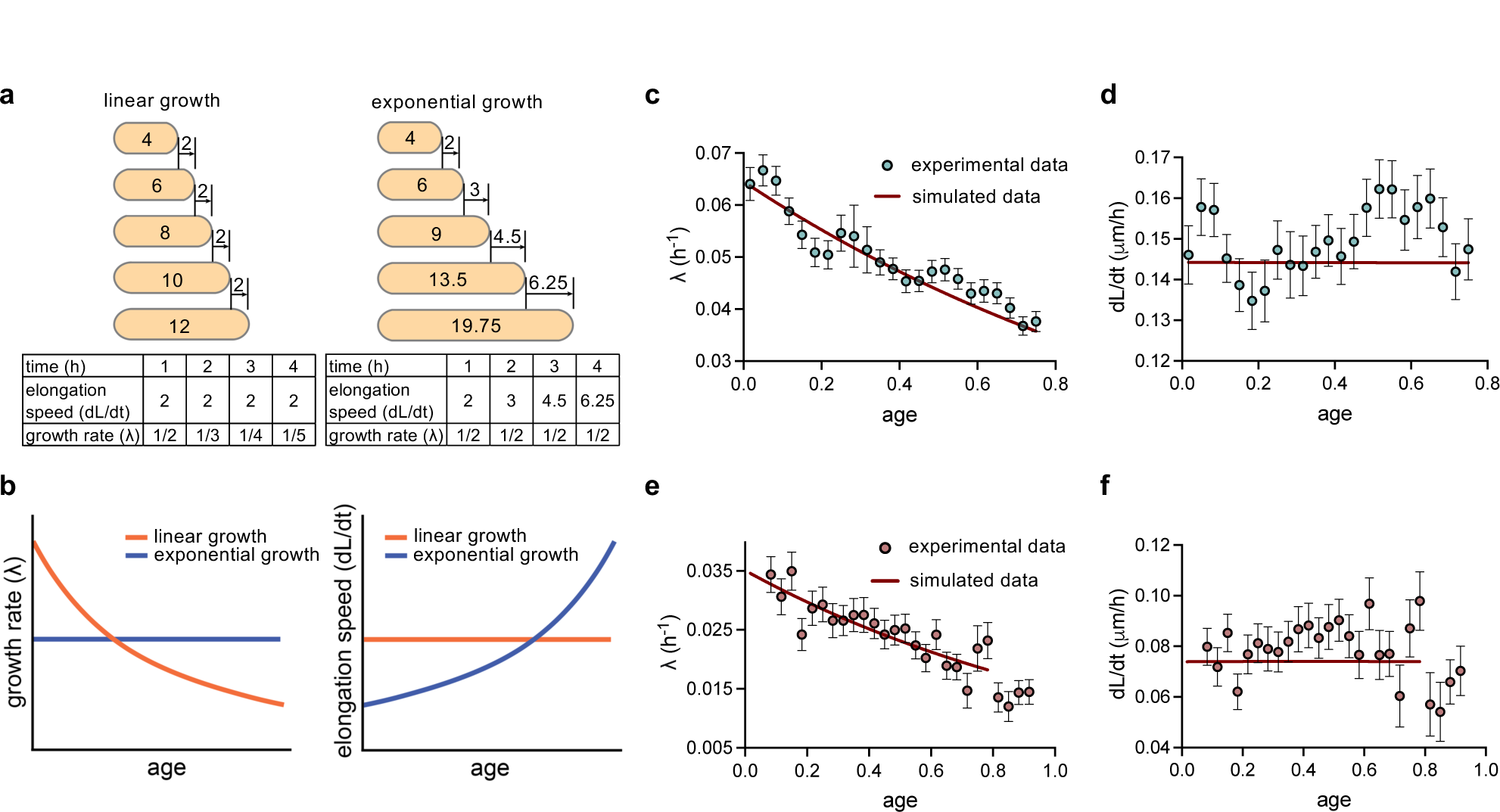
Linear and exponential growth modes. **a**, Schematic of length increase during linear growth (left side) and exponential growth (right side). The elongation speed is constant for linear growth, while for exponential growth, the growth rate 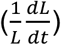 is constant. **b**, Predictions of growth rate vs. age and elongation speed vs. age plots for different growth modes. When plotted with growth rate vs. age, the binned data trend is a constant line for exponential growth, while for linear growth, the trend decreases with increasing age. For elongation speed vs. age, the binned data should increase throughout the age for exponential growth, while the trend is a constant line for linear growth. **c**, The binned data trend (blue dots) of growth rate 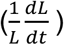 vs. age plot is shown for Mtb data (movie set A). The binned data trend decreases with age and is largely consistent with the predicted trend obtained using simulations of linear growth (red line). **d**, The binned data trend (blue dots) of elongation speed vs. age plot is shown for Mtb data (movie set A). The binned data trend is nearly constant, consistent with linear growth simulations (red line). **e**,**f**, The binned data trend (orange dots) of growth rate 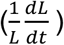 vs. age and elongation speed 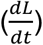 vs. age plot is shown for Mtb growing in an acidic condition (movie set C). The growth rate decreases with age (**e**) while the elongation speed is nearly constant (**f**), consistent with the predicted trend for linear growth obtained using simulations (red line).

For each cell, we calculated a growth symmetry metric, which is the proportion of total growth over one cell cycle that is contributed by the old pole. A growth symmetry score of 0.5 indicates symmetric growth (equal growth from the old and the new old), and scores <0.5 and >0.5 indicate more growth contribution by the new and the old poles, respectively. The distribution of growth symmetry is centered around 0.54 (Fig. 3c), indicating that the old pole elongates more than the new pole in a larger subpopulation of Mtb.

### Mtb growth is linear at the single-cell level

The similarity in the heterogeneity of cell cycle characteristics, such as cell size at birth and division between Mtb and *M. smegmatis,* made us question whether they grow in a similar manner, qualitatively, at the single-cell level. In previous studies, *M. smegmatis* has been observed to grow exponentially^24, 33^, similar to reports in *E. coli*^41, 42, 51–53, 55^. For exponentially growing cells, the increase in length per unit of time at each time point (elongation speed) is proportional to the length of the cell, with the proportionality constant equal to the growth rate (Fig. 4a,b). Another study reported biphasic growth of *M. smegmatis* cells^86^, and bilinear growth has also been proposed in *E. coli*^88^ where cells grow linearly for some time at an elongation speed after which they change to a different elongation speed. For linear growth, the elongation speed is constant (Fig. 4a,b). Thus, the mode of growth of *E. coli* has been a subject of debate. A previous study argues in favor of the use of a differential method for the analysis of growth during the cell cycle, i.e., plots of changes in cell length *dL/dt* vs. time since birth (*t*) are used instead of cell length (*L*) vs. *t*^89^. Recent works showed that differential methods such as elongation speed 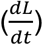 vs. age 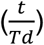 and growth rate 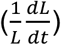 vs. age plots are appropriate methods for understanding the mode of growth^58, 60, 61^. The study found *E. coli* to grow super-exponentially (faster than exponential growth) in length^58^. Biphasic growth behavior in *B. subtilis* and an asymptotic linear growth pattern in *C. glutamicum* was observed as evidenced by growth rate vs. age plots and elongation speed vs. time plots^61, 62^. In this study, we also use the growth rate vs. age and elongation speed vs. age plots to elucidate the mode of Mtb growth.

For exponentially growing cells, the growth rate 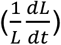 is constant throughout the cell cycle while the elongation speed 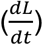 increases with age, as shown in the schematics in Fig. 4a,b. The best linear fit of the growth rate vs. age plot would be a horizontal line with a y-intercept equal to the average growth rate (Fig 4b). In contrast, for linear growth, the binned data trend for elongation speed will be constant throughout the cell cycle, and hence, the growth rate decreases with age (Fig. 4a,b)^58^.

On plotting the growth rate vs. age plot using data of Mtb grown in unbuffered medium (pH ∼6.8, n = 363 from movie set A, as described in Fig. 1), we find that the growth rate decreases with age (Fig. 4c) while the elongation speed vs. age plot stays constant (Fig. 4d). We found that the binned data trend from the experiments were largely consistent (barring small fluctuations) with the trend obtained using simulations of linear growth (Fig. 4c,d and Extended Data Fig. 6 and simulations in Methods). We also accounted for small changes in the binned data trends in the simulations due to measurement noise (Extended Data Fig. 6 and Model in Supplementary Information). The binned trends of elongation speed as a function of age show deviations of ∼25% from the ideal linear growth trend (red line in Fig. 4d) in simulations of linear growth with the same number of cell cycles as experiments (Extended Data Fig. 7). This matches the fluctuation levels in the binned data trend found in experiments (Fig. 4d). The mode of growth was also found to be linear when the experimental replicates were analyzed separately (Extended Data Fig. 5).

**Fig. 5.**
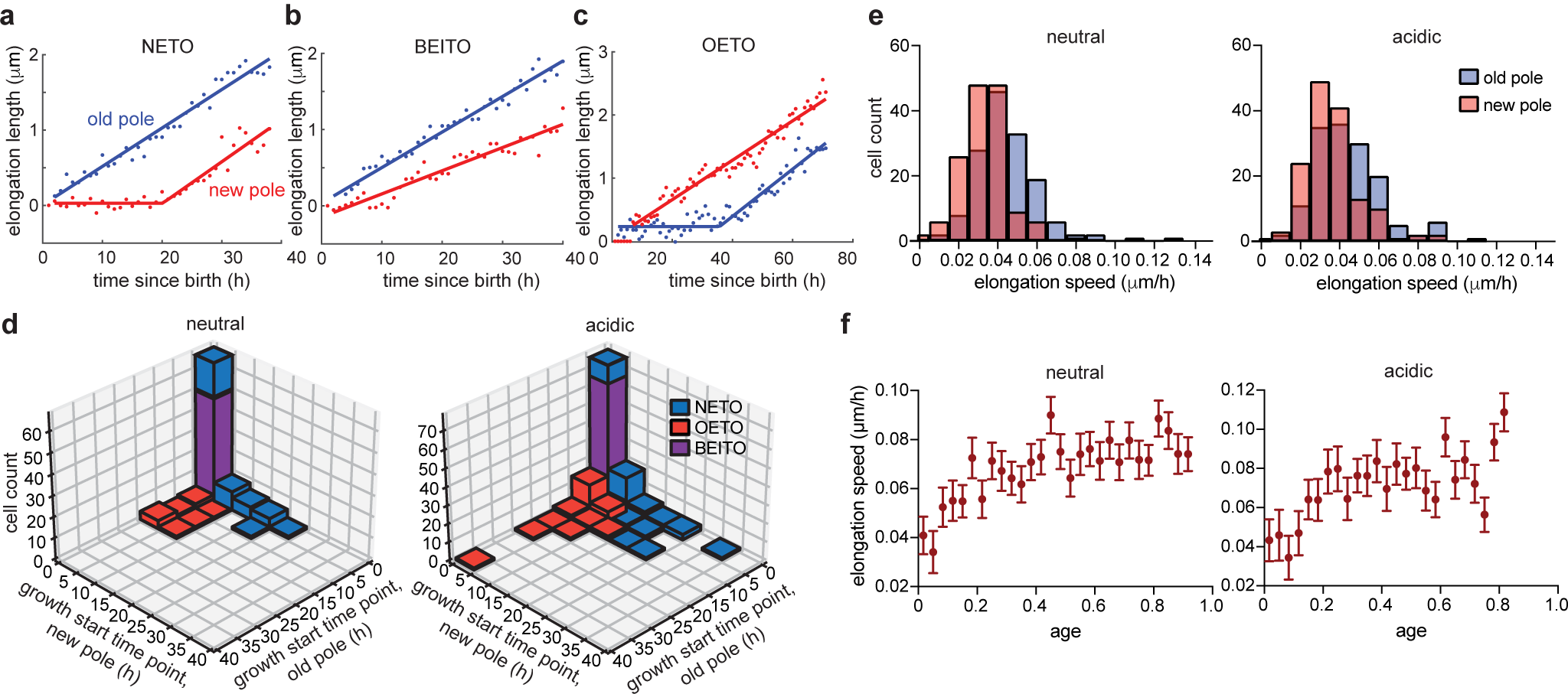
Growth characteristics at old and new poles. **a-c**, Representative examples of different polar growth dynamics (movie set B, other examples are shown in extended data fig. 4 and 10). the elongation length at each pole is shown as a function of time. The lines represent the best fit to the data and can be either linear or bilinear. **a**, The new pole starts elongating later than the old pole (new end take off, NETO). **b**, Both poles elongate from the beginning of the cell cycle (both ends immediately take off, BEITO). **c**, The new pole starts elongating before the old pole (old end take off, OETO). **d**, Joint distribution of the timings when the old and new poles start growing in neutral and acidic pH (movie set B and C, respectively). Blue, red, and purple bars indicate NETO, OETO, and BEITO cells, respectively. **e**, Distribution of the elongation speed at each pole in neutral and acidic pH (movie set B and C, respectively). **f**, Elongation speed vs. age curve obtained on simulation of a model where both poles could start growing from the beginning of the cell cycle or be delayed by a certain lag time.

### Mtb maintains a linear growth mode in an acidic medium

To understand whether the linear growth behavior of Mtb is specific to standard, rich growth conditions (unbuffered) used in this study, we also evaluated Mtb growth in an acidic medium. An acidic environment reflects a stressor that Mtb faces during infection, notably in phagolysosomal compartments in infected macrophages (pH 5.8∼6.2)^90, 91^. To measure growth mode in this slower-growing condition, we measured how Mtb grows in an acidic environment using time-lapse imaging (movie set C). By plotting the growth rate vs. age and elongation speed vs. age plots (pH 5.9, n = 135), we observed that the growth rate decreased with age while the elongation speed was mostly constant (Fig. 4e,f and Extended Data Fig. 8b). We observed that linear growth is also maintained when Mtb grows in a buffered neutral pH (pH 7.0) (Extended Data Fig. 8a and 9, movie set B). Overall, the binned data trends are largely consistent with linear growth for three different growth conditions (pH ∼6.8 with the SSB reporter in Fig. 4c-d (movie set A), pH 5.9 with FDAA pulse labeling in Fig. 4e-f (movie set C), and pH 7.0 with FDAA pulse labeling in Extended Data Fig. 9) (movie set B).

### Polar growth can start from either the old pole or the new pole or both at once

A previous study on polar growth in *M. smegmatis* indicated that Mtb elongate via NETO dynamics where the old pole starts growing linearly from birth while the new pole undergoes linear growth after a certain lag time^86^. This NETO dynamics predicts a growth change when the new pole starts growing. In this section, we investigate how to reconcile polar growth with the roughly linear growth we observed.

As introduced previously, pulse label experiments were used to observe growth at the old and new poles (Fig. 3a,b). We used mathematical modeling and *AIC* and *BIC* criteria to characterize the polar growth at each pole (Methods). Based on NETO, we expected the old pole to grow from birth and the new pole to be delayed before growing^86^ We show one example of this NETO dynamics with a trajectory of length grown vs. time in Fig. 5a and Extended Data Fig. 10a and 10d, where the green denotes the length grown by the old pole and the new pole growth is shown in red. However, as shown in Fig. 5b and Extended Data Fig. 10b and 10e, approximately half of the cells started growing from both poles within 1 h of cell birth (45.6% in neutral pH, 49.5% in acidic pH,, from n = 147 and 101 cells in movie sets B and C, respectively). Following the “NETO” nomenclature, we call these dynamics “BEITO” (both ends immediately take off). The joint distribution for the time (relative to birth) at which the old pole and the new pole start growing peaks at short times on both axes in both neutral and acidic pH (Fig. 5d). Unlike NETO dynamics^86^, we also find cases where the new pole starts growing before the old pole as shown in Fig. 5c and Extended Data Fig. 10c and 10f (“old end take-off”, OETO). In neutral pH, we found that NETO was observed in 33.3% of the total cells, while OETO was observed in 21.1% of the total cells. In acidic pH, the proportions for NETO and OETO were 37.6% and 12.9%, respectively (Fig. 5d). The elongation speeds are greater for the old pole growth as compared to the new pole growth in both neutral and acidic pH conditions (Fig. 5e). The overall greater elongation from the old pole relative to the new pole may be due to a combined effect of slightly more cases of NETO than OETO and a faster elongation speed from the old pole (Fig. 3d).

To reconcile the pattern of nearly consistent linear growth across the cell cycle (Fig. 4) and the pattern of polar growth with different ends growing at different times (Fig 5), we simulated growth mode behaviors using the polar growth data. In about half of the cells, both poles start growing simultaneously from the beginning of the cell cycle mimicking the BEITO dynamics. In instances where elongation does not commence at both poles upon cell birth, the time at which elongation begins at either of the poles is drawn from experimentally determined distributions, mimicking OETO and NETO (see Simulations in Methods section). Our simulations, using experimentally derived parameters from both acidic and neutral pH conditions, produced elongation speed vs. age trends that exhibited a similar qualitative pattern to the experimental data (Fig. 5f and Extended Data Fig. 8). Notably, a small increase in elongation speed at the start of the cell cycle was observed in both the simulations and experiments, resulting from NETO or OETO in approximately half of the cells.

To conclude, the cells undergo polar growth, with both poles starting to grow from the beginning of the cell cycle in half of the cells (BEITO). In the other half of the Mtb population, either of the poles can start growing first, in contrast to the NETO model, where the old pole always grows first. Using simulations based of our experimental data and distribution of cells that are BEITO, NETO, and OETO, we could reproduce the largely constant elongation speed vs. age curve observed in the experiments.

## Discussion

Using time-lapse imaging on Mtb reporter strains and Mtb labeled with cell-wall stains, we describe the growth behaviors and heterogeneity for tubercle bacilli in detail. The slow growth of Mtb may have led to exaggerated growth behaviors compared to *M. smegmatis*. However, prior studies using fixed-cell imaging suggested that polar growth in Mtb was less asymmetric overall than in *M. smegmatis*, which would attenuate differences in growth behaviors between sister cells and overall variation in growth behaviors^29, 86^. We show that cell-to-cell variation in cell size, interdivision time, and growth speed was similar and high compared to model rod-shaped bacteria. In *M. smegmatis*, a significant degree of variation in growth behaviors is correlated with the age of the growth pole^24, 26, 29^. However, there was no such correlation in Mtb, so sisters inheriting old growth poles are not much more likely to be longer and faster growing than their new pole-inheriting sisters.

Combining these single-cell growth data with mathematical modeling, we show that Mtb growth mode at the single-cell is linear throughout the cell cycle. Most bacterial species are thought to grow exponentially in biomass and volume, with *E. coli* recently reported to grow super-exponentially in length^42, 53, 54, 56, 58^. We found previously that *M. smegmatis* growth is exponential^24, 33^. *B. subtilis* grows in two distinct phases: linearly until cells reach a certain size, then exponentially until division^61^. A previous study on *C. glutamicum* proposed an asymptotically linear model where the bacilli grow exponentially and then grow linearly^62^. Messelink and colleagues suggested that asymptotically linear (elongation speed increases and then saturates) growth mode resulted from polar growth being the rate-limiting process. We showed that Mtb growth was nearly linear in slower, more stringent growth conditions in an acidic medium, mimicking a stressor the bacteria face during infection. The persistence of linear growth behavior in both rich and acidic conditions suggests that linear growth is the normal growth mode of Mtb and not the exception.

The linear growth mode of Mtb is achieved by linear growth from both poles, with the old pole elongating faster (Fig. 5). We found a distribution of biphasic growth behaviors in the population with both poles elongating from very early in the cell cycle in about half of the population (Fig. 5d). In the subpopulations where one pole begins to elongate before the other, we found that the first pole to elongate could be either the old pole or the new pole (Fig. 5d).

How bacteria grow at the single-cell level is a fundamental question in bacterial physiology. An important framework relies on the autocatalytic nature of ribosome production^38^. Denoting the fraction of ribosomes that are actively transcribing ribosomes by *ϕ*_*R*_, one finds that 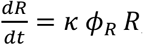, with *k* the translation rate. In balanced growth, the cellular growth rate must equal that of ribosomes, and the above equation implies a linear relation *λ* ∝ *ϕ*_*R*_, consistent with the linear dependency observed in *E. coli*^38^. Importantly, the above equation also predicts exponential growth at the single-cell level, inconsistent with our results for Mtb.

Linear growth at the single-cell level might arise due to several mechanisms. One possibility, natural for tip-growers such as Mtb, is that growth is limited by cell wall growth and that the amount of cellular machinery positioned at the tip is bounded – leading to a maximal tip velocity. Alternatively, the model in a previous study predicts that protein production is not limited by ribosomes for a sufficiently large volume/DNA ratio but rather by mRNA numbers^39^. However, mRNAs, in turn, are limited by the limited DNA content. This leads to linear growth at the single-cell level, consistent with experiments on perturbed budding yeast cells^92^. In the future, this hypothesis may be tested in Mtb using mutants where the volume/DNA is perturbed.

In addition to exploring the question of how linear growth is observed in Mtb, one may also inquire about the underlying reasons for this phenomenon. In bacteria such as *E. coli*, the existing paradigm is that the selection pressure faced by cells during evolution has led them to grow as fast as possible (barring the presence of non-growing subpopulations such as persisters^17, 93^. Indeed, in long-term-evolution-experiments growth rate has significantly increased over the generations^94^, and extensions of the ribosome-limited model for growth showed that by making ribosomes consist of numerous, smaller proteins leads to a significant growth-rate increase^95^, thus interpreting the ribosome structure itself in terms of growth rate optimization. This picture appears invalid in Mtb, suggesting it faces different constraints and selection pressure. This could arise due to its constant battle with the immune system, making its main challenge evading it rather than optimizing growth rate.

Slow growth may be a way in which the bacterial population survives the host immune response and establishes a chronic infection^96–100^. In the absence of fast growth and large population size, bet-hedging to survive stressors imposed on the bacteria during infection may be achieved by relatively high levels of cell-to-cell variation in a small population of cells^101, 102^. In contrast, fast-growing bacteria like *E. coli* reach large populations (and very quickly, with a growth rate >50x that of Mtb), so bet-hedging may be achieved by less variation but much higher cell numbers from faster growth^102^. As an example of bet-hedging, Basan et al. demonstrated that a trade-off between growth and survival (adaptation) in the fluctuating environment was made by the nutrient shift in *E. coli*. They showed that the *E. coli* population that invests more resources in growth experiences slow growth while adapting better when environmental conditions change^102^. Therefore, we speculate that the objective of growth behavior in Mtb may be fundamentally different from more well-studied bacteria; for example, Mtb grows more slowly with increased variation, and model bacterial species grow more rapidly with a more modest variation. We have yet to understand how linear growth enables Mtb to achieve an optimized growth rate and population-level heterogeneity. It may be that linear growth enables Mtb to adapt to the constantly changing microenvironment of TB granulomas during infection while optimizing cell state variation and population size. Alternatively, the linear growth mode might be a spandrel^103, 104^.

Our work provides a new baseline understanding of Mtb growth properties and their variations in a population of cells. These data include cell sizes, cell cycle timing, and elongation speeds. We find that the growth mode and level of heterogeneity are together unique and cannot be explained using models developed in model bacteria or other closely related species such as *M. smegmatis*. Given the critical role of growth, metabolic state, and adaptation to fluctuating environments that Mtb faces during infection to Mtb virulence and drug response, further studies on the growth behaviors of Mtb and other pathogens will enable us to develop improved therapeutic interventions.

## Acknowledgments

We thank S. Tan for the generous gift of the CDC1551 SSB-GFP, *smyc’*::mCherry strain. This work was funded by the NIH (R01 AI143611-01) to B.A. and in part by NSF CAREER (1752024) to A.A.

## Author contributions

E.S.C, P.K., B.B.A., and A.A. conceptualized the project, E.S.C. and M.K. conducted the experiments and time-lapse imaging annotation, and P.K. ran mathematical analysis. E.S.C., P.K., B.B.A., and A.A. wrote the draft, and all authors reviewed and edited the manuscript. B.B.A. and A.A. acquired funding.

## Methods

### Overview of the movie sets

For the baseline movie set (movie set A), we used Mtb CDC1551 strain harboring the SSB-GFP reporter, which was grown in unbuffered (pH ∼6.8) standard (supplemented 7H9) growth medium at 37°C. Time-lapse imaging was conducted for 96 hours, with images taken every hour.

Two additional movie sets were made using the FDAA pulse-labeled CDC1551 wild-type strain. Movies were made in neutral (pH 7.0, movie set B) or acidic (pH 5.9, movie set C) standard (supplemented 7H9) growth medium at 37°C. Time-lapse imaging was conducted for 140 hours, with images captured every hour.

### Mtb strain

Mtb CDC1551 strain (movie sets B and C) and its transformant with a hygromycin-resistant replicating plasmid expressing single-stranded binding protein fused to a green fluorescent protein (SSB-GFP) and *smyc’*::mcherry (movie set A) are used in this study^77^.

### Bacterial culture

Mtb was grown in a standard medium consisting of 7H9 broth (ThermoFisher; DF0713-17-9) with 0.05 % Tween 80 (ThermoFisher; BP338-500), 0.2 % glycerol (ThermoFisher; G33-1), 10% Middlebrook OADC (ThermoFisher; B12351). For the SSG-GFP reporter strain (movie set A), 50 µg/ml of hygromycin (ThermoFisher; 10687010) was added to maintain SSB-GFP, *smyc’*::mcherry. Mtb strain was grown to OD_600_ of 0.5 - 1.0 from frozen aliquots at 37°C with mild agitation. Cultures were subcultured via back dilution to OD_600_ 0.05 and grown to the mid-log phase (OD_600_ 0.5-0.7) before experimental use. For movies where polar growth is assessed (movie sets B and C), Mtb cells were backdiluted to OD_600_ of 0.2 in 10 ml 7H9 media (unbuffered) supplemented with 100 µm HADA (see below) for 24 h. Labelled cells were washed twice with PBS (ThermoFisher; 20012-027) + 0.2 % Tween-80 (PBST) and resuspended in pH 5.9 or 7.0-adjusted fresh 7H9 media that was supplemented with sterile spent medium (50:50) for loading into the microfluidic devices for time-lapse imaging.

### FDAA labeling (movie sets B and C)

The blue fluorescent D-amino acid (FDAA), HADA (Tocris; 6647), was used for movie set B and C. HADA powder was dissolved in DMSO to a stock concentration of 100 mM and stored short-term at −80 °C. Cells were incubated in 100 µM HADA for 24 h prior to the start of the imaging.

### Live-cell microscopy

Before loading Mtb cells into a custom polydimethylsiloxane (PDMS) microfluidic device, cells were filtered through a 10 μM membrane filter to remove clumps. Mtb cells were loaded into a microfluidic device, as previously described (Richardson et al., 2016)^22^. The devices contain a main microfluidic feeding channel with a height of 10-17 μm and viewing chambers with a diameter of 60 μm and a height of 0.8-0.9 μm. Fresh medium was delivered to cells at 5 μl/min flow using a microfluidic syringe pump. The device was placed on an automated microscope stage within an environmental chamber maintained at 37 °C. Mtb cells were imaged using a widefield Deltavision PersonalDV (Applied Precision, Inc.). For movie set A, cells were illuminated with an InsightSSI Solid State Illumination system every hour for 96 hours. Cells were imaged using transmitted light brightfield microscopy, GFP (475 nm/525 nm), and mCherry (575 nm/625 nm). mCherry was imaged to ensure the presence of the plasmid and was not used for analysis. The movies were performed in biological triplicate, with each replicate performed separately in different microfluidic devices on different days. For FDAA pulse-labeled movies (movie sets B and C), cells were imaged every hour for 140 hours using transmitted light brightfield microscopy, and HADA was visualized with CFP filter (433 nm/475 nm).

### Live-cell image segmentation

For movie set A, ImageJ plugin ObjectJ was used to hand-annotate cell length, growth, and cell cycle progression throughout the image stack (time-lapse). The SSB-GFP reporter forms a green fluorescent focus during DNA replication. Newly divided cells generally did not have SSB-GFP focus, labeled as the pre-replication period (B period). The period where one or two SSB-GFP foci were detected was defined as the replication period (C period), and the subsequent period where foci were absent was labeled as the post-replication period (D period). Some cells had one or two foci after the division period but before septation; in these cases, we labeled those occurrences as the pre-division period (E period)^24, 77, 78^. Cell poles and visible foci were annotated in each frame - two points at each pole of a single cell were annotated if no foci were detected, while three (one focus) or four (two foci) points were annotated when foci appeared. The localizations of foci were manually analyzed (custom code) and used to determine cell cycle timing. The annotation in each frame was extracted, containing information on cell length and cell cycle progression over time (1-hour time scale). Information on mother-daughter and accelerator-alternator cell relationships was also collected from cell pedigree trees. During time-lapse imaging, we observed a rare subpopulation in which the cells express a high intensity of GFP. We observed that these cells did not divide and entered growth arrest. This may be due to phototoxicity caused by high SSB-GFP abundance. These cells were excluded from annotation.

For FDAA pulse-labeled movies (movie sets B and C), cell poles and HADA labeling were hand-annotated in each frame using ImageJ (version 1.53f) with an ObjectJ plug-in. Whole-cell labeling was annotated at each pole in the first time point. When cells elongated, a non-labeled area appeared, representing the new growth site. In cases where cells elongated from only one pole, three points were annotated, starting from the newly growing pole (unlabeled) to the HADA-labeled area which includes the other pole. When cells elongated from both poles, four points were annotated, starting from the newly growing pole (single point), the HADA-labeled cell body (two points), and the other pole (single point). After annotation, the x and y coordinates of each annotation point were extracted, and the distance was calculated using the Euclidean distance formula.

### Statistical analysis

The scatter plots are presented with median values. Mann-Whitney test was performed throughout the features compared between accelerator and alternator cells within Mtb or *M. smegmatis* strains. Significance between CV values was tested using an asymptotic test for the quality of coefficients of variation from k populations^105^. *P* values < 0.05 were considered statistically significant.

### Simulations

In the main text, we showed Mtb were largely consistent with linear growth. We used growth rate vs. age and elongation speed vs. age plots to reach the conclusion. Before we could interpret the experimental results using these methods, we wanted to test the methods against model simulations. The data analysis used an underlying model of linearly growing cells with measurement noise. Simulations of linearly growing single cells were carried out over 1000 generations. Elongation speeds were assumed to have a Gaussian distribution with mean ⟨*λ*_()*_⟩ and coefficient of variation *CV*_*λ,lin*_ determined using the experiments. The elongation speed was determined at the start of the cell cycle and was assumed to be independent of the elongation speed of the previous cell cycle. The cells divided upon reaching size *Ld* = 2(1 − *α*)*Lb* + 2*α*Δ + *η*. Here, Δ is a constant and *α* is the size regulation strategy which can take any value from 0 to 1. When *α* is 1, cells divide on reaching a particular size 2Δ (sizer) and for *α* = 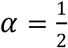, cells divide on adding a constant size Δ from birth. *α* and Δ are determined using the slope and intercept of the experimental *Ld* vs. *Lb* plot which are 2(1 − *α*) and 2*α*Δ, respectively (Amir, 2014). *η* is the size additive division noise with mean zero and standard deviation set such that the CV of *Lb* in simulations matched that of experiments. Our results are independent of the nature of division noise. Upon division, the cells divided symmetrically on average with the division ratio *r* = 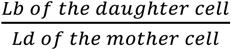 drawn from a Gaussian distribution with mean = 0.5 and standard deviation, *σ_r_* ≈ 0.06 (determined using mother-daughter pairs in experiments). The length of each cell is measured at 1-h intervals. The measured length is the sum of the actual length of the growing cell and a measurement error normally distributed with mean zero and standard deviation determined from experiments (see Supplementary Information).

We also investigate the effects of NETO, OETO and BEITO dynamics on the binned data trends of growth rate vs age and elongation speed vs. age plots. We hypothesize a model where roughly half of the cells follow BEITO dynamics. For these cells, it is assumed that the elongation speeds of the poles are independent of each other and drawn from a normal distribution at the beginning of each cell cycle. The mean and standard deviation of this distribution are determined from the experimental data. As mentioned in the main text, for the rest 50% cases with OETO and NETO dynamics, the time at which elongation begins at either of the poles is drawn from experimentally determined distributions. The cell grows linearly at each of the poles and divides when the cell length reaches a length *Ld* which depends on the cell size at birth *Lb*, *Ld* = 2(1 − *α*)*Lb* + 2*α*Δ + *η* (similar to linear growth simulations above). *η* is a Gaussian size additive division size noise with mean 0 and standard deviation fixed such that the standard deviation of the Lb from the simulations matches that of experiments. The nature of the noise (time additive or size additive) does not change the qualitative nature of the elongation speed vs. age plots. The cell divides symmetrically on average, with the standard deviation of division asymmetry determined from experiments. The model is simulated for a single lineage of n generations (n = 100 for acidic pH and n = 147 for neutral pH). The length of the cell is stored at 1 h intervals mimicking the experiment. The recorded cell length is the sum of the actual length of the cell and a noise due to measurement error (see text in Supplementary Information). We ignore cells where one of the poles does not grow during the cell cycle.

### Characterizing single trajectories using AIC and BIC values

We characterize the growth at the old and new poles using FDAA (HADA) labeling (Fig. 3). Fig. 3a (middle panel) shows that the cells have an unlabeled part at the old pole after the first generation. The old pole growth is measured by adding unlabeled parts to the existing unlabeled region. The growth at the new pole is marked by the appearance of an unlabeled region at the new pole and, in a few cases, the growth of unlabeled parts to the existing unlabeled region. The aim is to identify the amount of growth at each pole and the time at which the pole growth starts. However, precise measurement of single-cell polar growth is difficult because the clumping of Mtb cells (Extended data Fig. 1a) obscures the position of the poles and, thus, complicates the determination of the HADA unlabeled part. This section explains the statistical models used to determine the polar growth at both ends at a single-cell level.

Previous studies have determined growth at each individual pole to be linear^86^. Linear growth is also supported by our results in the main text (Fig. 4). Thus, to determine the growth at each pole, we fit two different models to the length vs. time trajectories – 1. linear growth 2. bilinear growth is where the length stays constant for a certain time and then increases linearly. The bilinear growth was used to characterize the NETO^86^, where it was proposed that the new pole starts growing after a time delay from cell birth. We assume that some old poles may also grow bilinearly (Fig. 5c, Extended Data Fig. 4a, and Extended Data Fig. 10c and 10f).

In cells that already have an unlabeled part at the old pole, we calculate the amount of old pole growth at time *t* from birth as the difference between the length of the unlabeled HADA region at the old pole at time *t* from birth and the initial unlabeled part at birth. The measured length grown can be negative for the old pole as the unlabeled HADA region at birth can be inaccurate due to cell clumping or cell tilting along the z-plane in the microfluidic chamber. We show examples of length grown vs. time for the old pole (blue) and new pole (red) in Fig. 5, Extended Data Fig. 4 and Extended Data Fig. 10. In most of the cells of the pulse label experiment, we do not observe a HADA unlabeled region at the new pole at the time of birth. In these cells, the length grown at the new pole is the length of the unlabeled HADA region at the new pole. The length grown is marked as zero when we do not observe the unlabeled HADA region at a pole.

Next, we fit the two models discussed above for each single cell to the length grown at each pole vs. time curves. The linear model is characterized by two parameters *y* = *ax* + *b* where *a* is the elongation speed of the pole, and *b* is a measure of the unlabeled HADA region at cell birth. For the bilinear model, the underlying equation is,

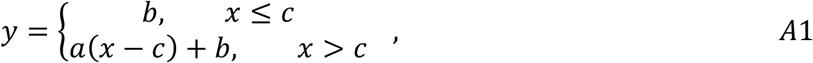

where *a*, *b*, and *c* are the elongation speed of the pole, bias in determining the initial HADA unlabeled region and time when the pole starts growing (relative to birth), respectively. For fitting, we ignore those data points where the length grown is zero (the y-axis is zero). We minimize the squared sum of residuals (*RSS*); 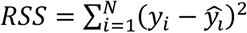, where *y*_*i*_ is the true value of the *i*^*th*^ data point of the dependent variable, and *ŷ*_*i*_ is the predicted value from the model. To compare which of the two models better fits the single-cell trajectories, we use the Akaike information criterion (*AIC*) and the Bayesian information criterion (*BIC*). They are defined as,

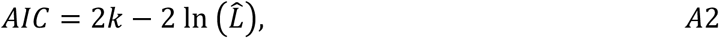

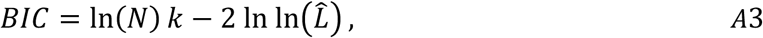

where *k* is the number of parameters in the model (*k* = 2 for linear, *k* = 3 for bilinear), *N* is the number of data points fitted and *L* is the maximum value of the model’s likelihood function. In both *AIC* and *BIC* methods, a model is favored if it has a greater maximum likelihood value, and it is penalized for having a greater number of parameters, the penalty being different for *AIC* and *BIC* as shown in Equations *A2* and *A3*. Assuming that the errors (*y_i_* − *ŷ_i_*) are drawn from an independent and identical normal distribution, the *AIC* and *BIC* values can be simplified to^106^,

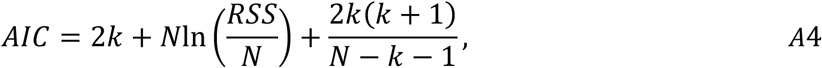

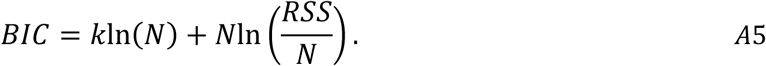

The *AIC* and *BIC* values themselves have little significance. The relevant metric is the difference between *AIC* and *BIC* values of the two models being compared, in our case, they are *ΔAIC* = *AIC*(*linear*) − *AIC*(*bilinear*), *ΔBIC* = *BIC*(*linear*) − *BIC*(*bilinear*) . A model with a lower *AIC* and *BIC* value is preferred. In our case, the bilinear model is preferred if both *ΔAIC* and *ΔBIC* values are greater than zero, while linear is preferred if the *ΔAIC* and *ΔBIC* values are less than zero. In a few cases where *ΔAIC* and *ΔBIC* values have opposite signs, the model with fewer parameters, i.e., the linear model is preferred.

Having chosen the appropriate model using the *ΔAIC* and *ΔBIC* values obtained upon fitting both models to the length vs. time trajectories of each cell, we aim to calculate the time at which the growth at a particular pole starts, the elongation speed at that pole, and the amount of growth at that pole. We discuss the calculation of these quantities for different cases of fit results obtained-first for old pole growth and subsequently for new pole growth.

A bilinear curve best fits the trajectory for old pole growth shown in Extended Data Fig. 4a. In these cases, where the old pole growth is bilinear, the value of parameter *c* (Equation *A1*) of the best bilinear fit denotes the time at which the pole starts elongating. The slope *a* is the elongation speed of the pole and the constant *b* is the bias in determining the initial unlabeled HADA region. Thus, the amount of polar growth during the cell cycle is the length increase within time *c* and *Td* (the doubling time). Mathematically, the amount of polar growth = *a*(*Td* − *c*). For the old pole growth trajectories in Extended Data Fig. 4b and c, where the best fits are linear, we assume that the growth starts from time 0. The y-intercept obtained (positive in (b) and negative in (c)) can be interpreted as the error in determining the initial unlabeled HADA region. The elongation speed is the slope of the best linear fit and the amount of growth is equal to *Slope* × *Td*.

Next, we discuss the calculations of growth parameters for the new pole growth. We fit the non-zero length grown vs. time data to the two models-linear and bilinear- and choose the appropriate model based on *ΔAIC* and *ΔBIC* values. In the trajectory shown in Extended Data Fig. 4a, the best fit is linear with a negative y-intercept. On extrapolating the best fit to y = 0, we obtain a positive time *T_trans_*. Since we do not observe the unlabeled HADA region before this time point, we interpret the time *T_trans_* as the time when the pole starts growing. The raw data shows the new pole to have HADA label for some time after *T_trans_* because the unlabeled HADA region is small and might be undetectable in the movies until it reaches a particular length. Examining the movies shows that this length is around 0.2-0.4 µm. This can also be seen in all the Extended Data Fig. 4 trajectories, where there is a sudden jump in the length of the new pole. The elongation speed is the slope of the best linear fit and the amount of growth is *Slope* × (*Td* − *T*_!"#$%_). In Extended Data Fig. 4b, the best linear fit has a positive y-intercept. In this case, we interpret the y-intercept as an initial undetectable HADA unlabeled region at the new pole. The new pole starts growing from the beginning of the cell cycle with an elongation speed equaling the slope of the best linear fit. The amount of growth at the new pole during a cell cycle is *Slope* × *Td*. In Extended Data Fig. 4c, where a bilinear fit better explains the length grown as a function of time, the interpretation of the fit is similar to that of the old pole. The constant parameter *b* in Equation *A1* denotes the HADA unlabeled region at cell birth, *c* denotes the time when the new pole starts growing and the slope *a* denotes the elongation speed. The amount of growth is given by the same equation as the old pole, *a*(*Td* − *c*).

In a few cases, we observed that the new pole had an unlabeled HADA region at cell birth. In such cases, we analyzed the new pole using the same method as the old pole, as discussed previously in this section.

## Code availability

Custom MATLAB code used in this study are available at https://github.com/pkar96/Mtb-growth

---

**Extended Data Fig. 1 Challenges of time-lapse imaging and annotation of Mtb.** (**A**) Clumps form as Mtb cells grow and divide (movie set A). The clumps make single cells hard to distinguish, preventing auto-segmentation. (**B**) V-snapping during cell division and formation of curved shapes (movie set A). The irregular cell shape and stickiness of tubercle bacilli prevents use of thin channel-based microfluidic devices. (**C**) Polar growth of Mtb cells (movie sets B and C). Each single cell was fully stained using a pulse-label of FDAA (green) before starting time-lapse imaging. Unlabeled regions of the cell wall indicate regions that elongated in the chase period during time-lapse imaging. Scale bar, 2 µm.

**Extended Data Fig. 2 Single-cell traces of SSB-GFP localization (movie set A).** The distance between the x-axis and the black circle indicates cell length. Yellow circles indicate SSB-GFP foci. (**A**-**C**) Cells with a cell cycle comprised of B-C-D periods. The duration of each period can vary between single cells. (**D**,**E**) Cells with cell cycle comprised of B-C-D-E periods. Cells with an E period initiate another round of DNA replication before division. The daughter cells of this type of cell lack the B period and enters C directly (see figure F). (**F**) Example cell with a cell cycle comprising a C and D period. This type of cell cycle occurs in cells whose mother begins replication before division (E period). In cases where foci disappeared for a frame or two due to focus issues and then reappeared, we assumed replication continued until the last foci disappeared.

**Extended Data Fig. 3 Growth characteristics of Mtb during the course of time-lapse imaging (movie set A).** Growth behaviors of length at birth, length at division, and the interdivision time for individual cells vs the time of their birth during the course of the movie. Different colors (gray, green, and orange) indicate biological replicates. Linear regression was used to determine slopes of Lb, Ld, and Td of each biological replicate. Each slope was not significantly different from zero (average p-value of biological triplicates: Lb 0.41, Ld 0.44, Td 0.35).

**Extended Data Fig. 4 Growth characteristics at old and new poles (movie set B).** Length grown at the old pole (green dots) and the new pole (red dots) is shown as a function of time for three different cells. Also shown are the best fits (green line for the old pole and red for the new pole) to the length growth vs time trajectories. A bilinear fit is the best fit for the old pole growth of cell (**A**) while the linear fit is preferred for the cells in (**B**) and (**C**) based on the AIC and BIC values. For the new pole growth, linear fits are preferred for cells (**A**) and (**B**) while a bilinear fit is preferred for cell (**C**).

**Extended Data Fig. 5 Growth rate vs age and elongation speed vs age plots of each biological replicate (movie set A).** (**A**-**C**) The binned data trend (blue dots) of growth rate 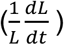 vs. age plots are shown for three individual replicates of Mtb data. The binned data trend decreases with age and is largely consistent with the predicted trend obtained using simulations of linear growth (red line). (**D**-**F**) The binned data trend (blue dots) of elongation speed vs. age plot is shown for the three replicates of Mtb data. The binned data trend is constant consistent with the predicted trend obtained using linear growth simulations.

**Extended Data Fig. 6 Binned data trend in growth rate vs age and elongation speed vs age plots.** (**A**,**B**) Variance of difference in lengths at time *kΔt* from birth (where Δ*t* =1 h) and the length at birth is plotted against time *k*Δ*t* (blue dots). The red line is the best quadratic fit of the form: *y*(*x*) = *ax*^2^ + *b*. The variance is calculated over all cells growing in acidic medium in pulse label experiments (**A**, movie set C) and unbuffered media with SSB-GFP label (**B**, movie set A). (**C**,**D**) Simulations of linear growth are carried out with the division size being determined by the birth size. For growth rate vs age (**C**) and elongation speed vs age (**D**) plots, we find the binned data trend for a single simulation run (blue dots). We also show the average binned data trend where the binned data trend for each of the plots are averaged over 1000 simulation runs. The solid red line represents the average binned data trend for simulations where the measurement error is non-zero (equal to that obtained from unbuffered Mtb movies) and the dashed red line corresponds to simulations with zero measurement error.

**Extended Data Fig. 7 Simulation runs of unbuffered Mtb movie.** (**A**,**B**) Simulations of linear growth are carried out with the division size being determined by the birth size. The parameters used in the simulations are determined from the unbuffered Mtb movies. The number of cell cycles considered (*n*) = 363 which is the same number of cell cycles in the Mtb experiments in unbuffered media. We show the binned data trend of elongation speed vs age (red dots) for two simulation runs.

**Extended Data Fig. 8 Simulation run of neutral and acidic elongation speed vs. age.** (**A**,**B**) The binned data trend of elongation speed vs age obtained from a single simulation run with parameters obtained from pulse label experiments in neutral medium (**A**) and acidic medium (**B**). The simulations take into account BEITO (both end immediately take-off), NETO (new end take-off) and OETO (old end take-off) dynamics. The division size of the cells depends on the birth size. The model is simulated for number of cell cycles (*n*) = 147 in (**A**) and 100 in (**B**).

**Extended Data Fig. 9 Growth rate vs age and elongation speed vs age plots of pH 7 condition (movie set B).** (**A**,**B**) The binned data trend (dots) of growth rate 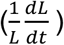 vs age and elongation speed 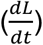 vs age plot is shown for Mtb growing in neutral pH conditions. The growth rate decreases with age (**A**) while the elongation speed is nearly constant (**B**) and it agrees with the predicted trend for linear growth simulations (red line) to a great extent.

**Extended Data Fig. 10 Single cell growth in NETO, BEITO, and OETO (movie set B).** The snapshots display the length grown at the old pole (green line) and the new pole (red line). The plots next to the snapshots illustrate the best fits for length growth vs. time trajectories. A linear fit best describes the old pole growth of cells (**A**) and (**B**), while a bilinear fit is preferred for the cell in (**C**) based on the Δ*AIC* and Δ*BIC* values. Regarding the new pole growth, a linear fit is preferred for cells (**B**) and (**C**), while a bilinear fit is preferred for cell (**A**). Specifically, (**A**) and (**D**) represent single cells with NETO, (**B**) and (**E**) represent single cells with BEITO, and (**C**) and (**F**) represent single cells with OETO.

