## Supplementary figures and images for "*Mycobacterium tuberculosis* grows linearly at the single-cell level with larger variability than model organisms"

### Extended Data Figure 1

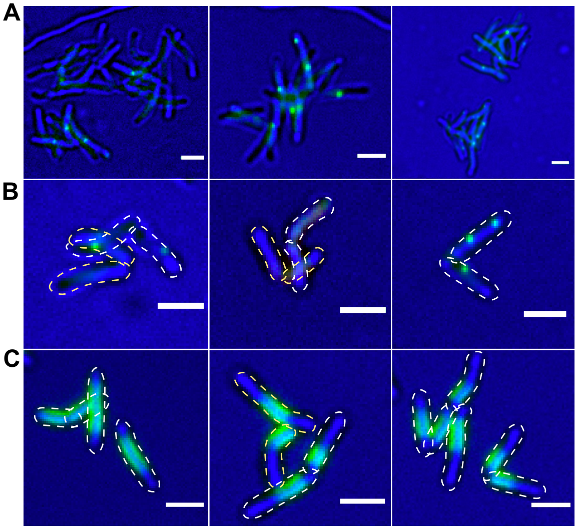

### Extended Data Figure 2

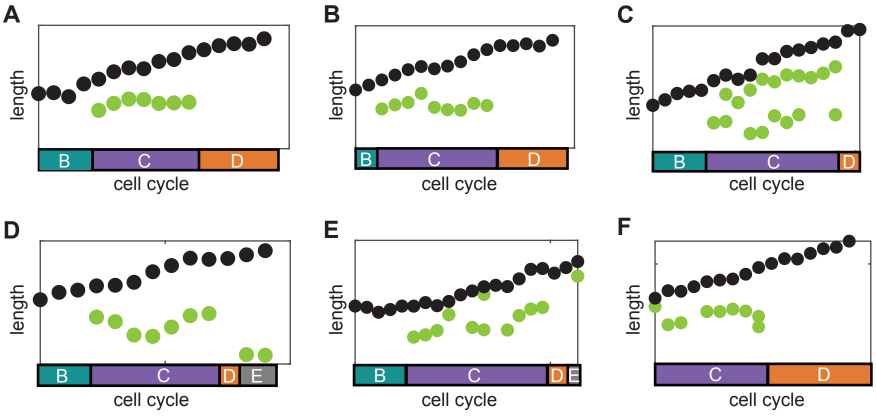

### Extended Data Figure 3

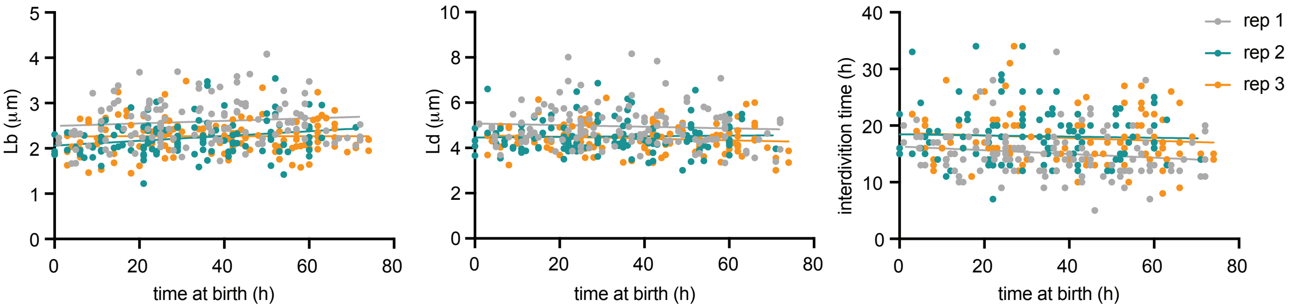

### Extended Data Figure 4

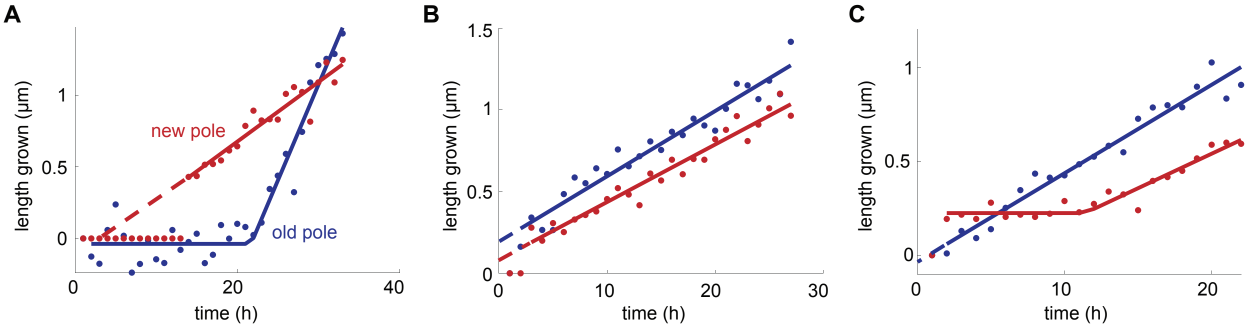

### Extended Data Figure 5

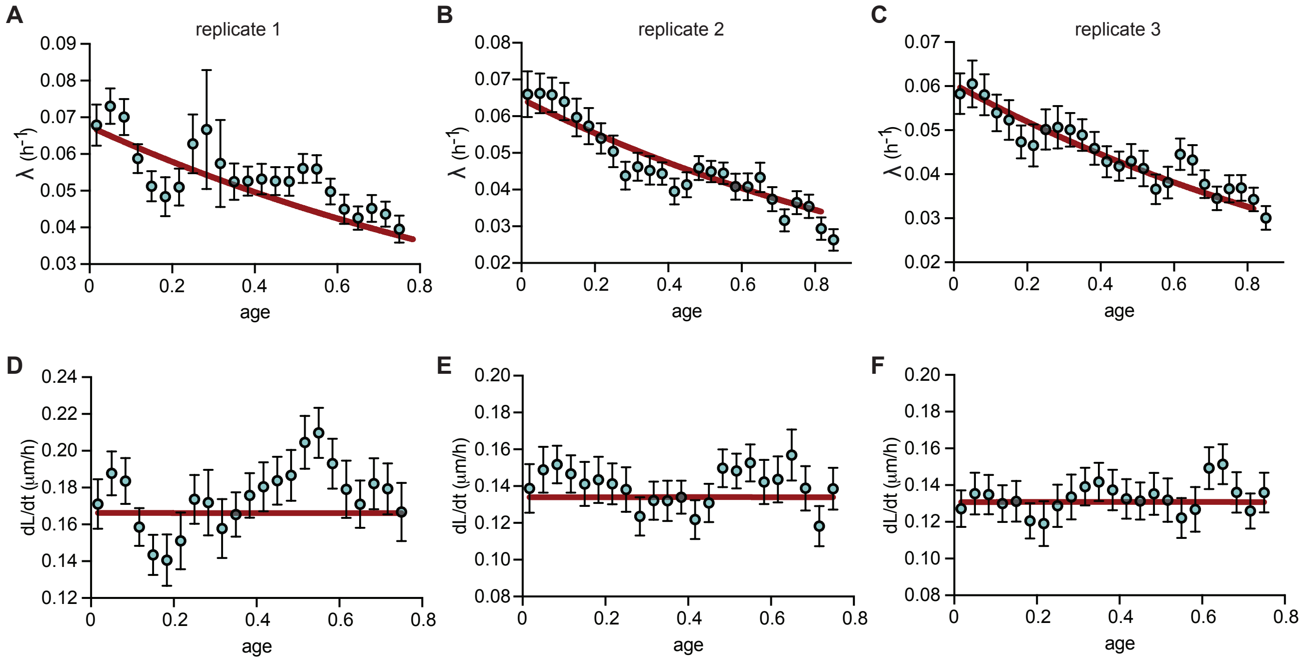

### Extended Data Figure 6

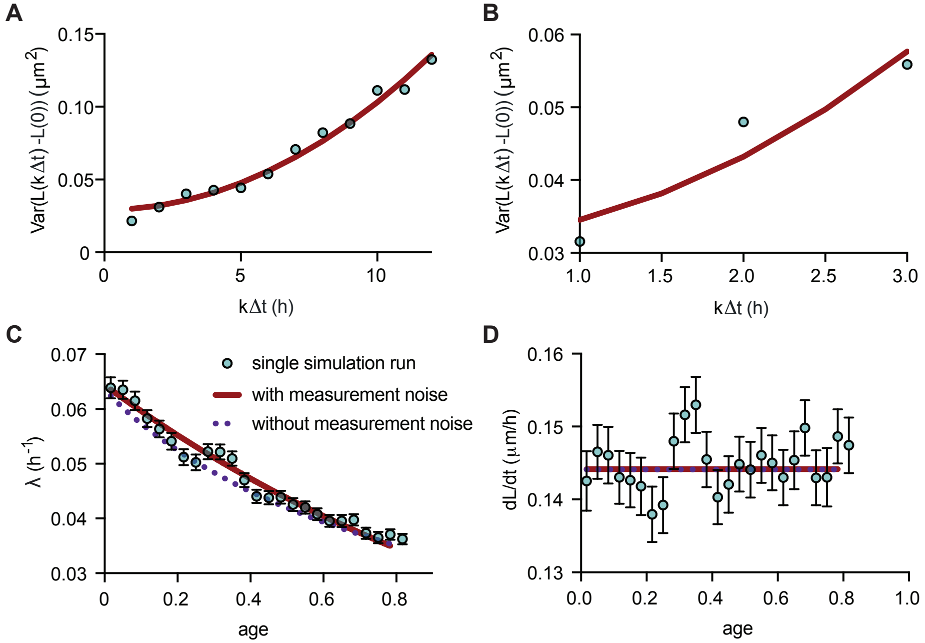

### Extended Data Figure 7

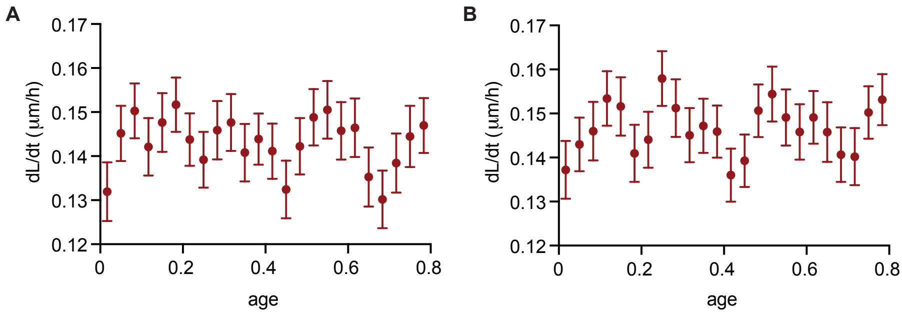

### Extended Data Figure 8

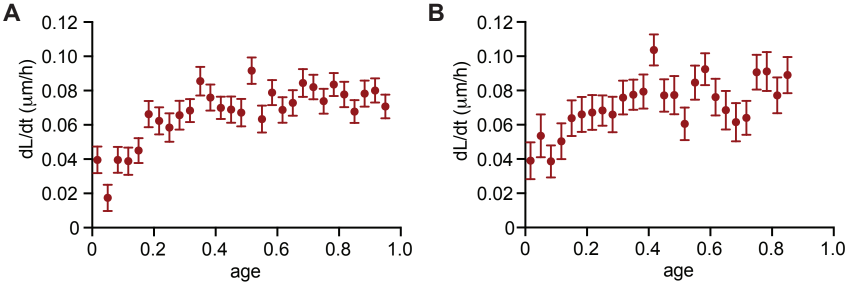

### Extended Data Figure 9

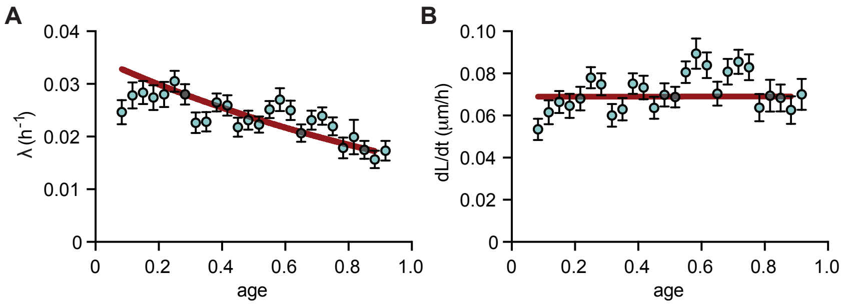

### Extended Data Figure 10

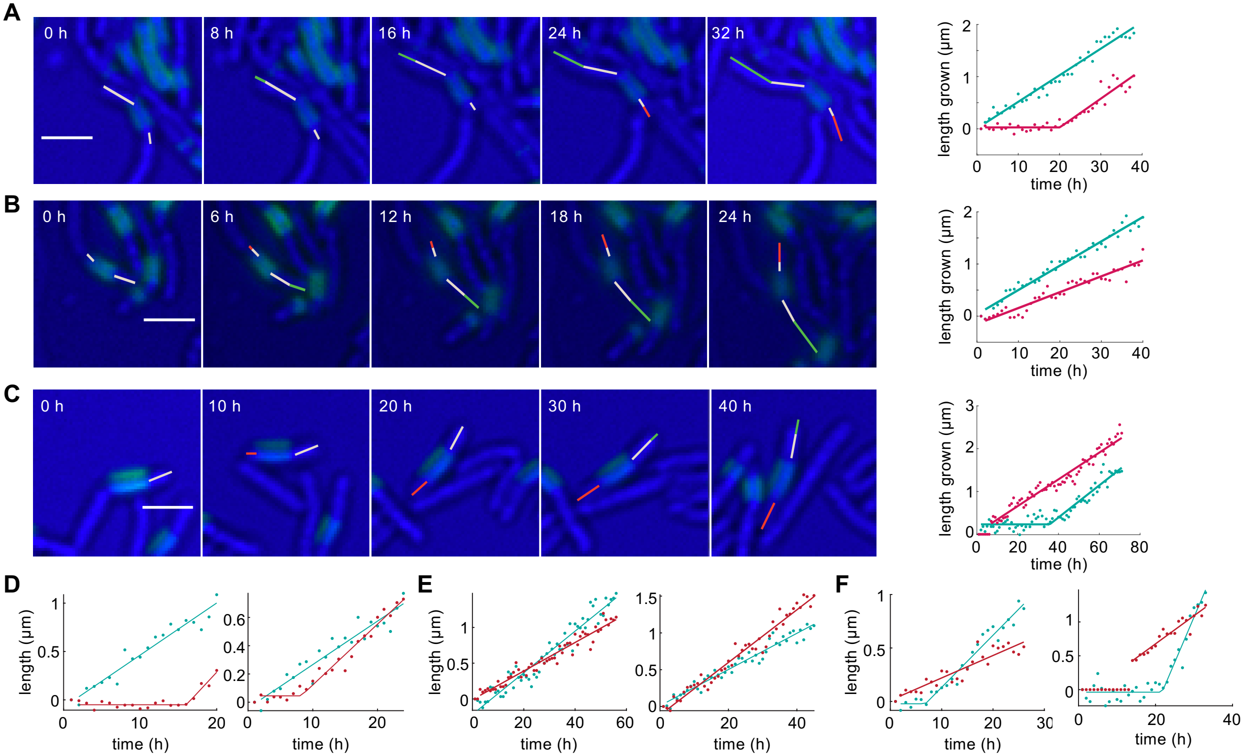
