## Supplementary Information for "*Mycobacterium tuberculosis* grows linearly at the single-cell level with larger variability than model organisms"

We show in the main text that the elongation speed vs. age and the growth rate vs. age plots point to the mode of growth being linear in Mtb. Here, we show that measurement errors in cell lengths have a small effect on the binned data trend in growth rate vs. age and elongation speed vs. age plots.

First, we will introduce a model based on which we will predict the binned data trend in growth rate vs. age and elongation speed vs. age plots.

**Model**

Assuming linear growth, the cells of length $L_a$ at time $t$ from cell birth grow as,

$$\frac{dL_a}{dt} = \lambda_{lin}. \qquad S1$$

In the model, $\lambda_{lin}$ is assumed to be constant within the cell cycle, with its value set at the time of birth. However, its value can vary between cell cycles with cell cycles averaged mean $\langle\lambda_{lin}\rangle$ and coefficient of variation $CV_{\lambda,lin}$.

Next, we will introduce measurement errors into the model, whose value will be estimated from our experiments. We assume that cells grow linearly according to Equation $S1$. The measured length $L(t)$ at time $t$ from cell birth varies from the actual length, $L_a(t)$ of the cell by a measurement error term, $\xi_m(t)$. Instrument error limits and imprecision during cell image segmentation can contribute to measured lengths being different from actual lengths. Thus, measured lengths are,

$$L(t) = L_a(t) + \xi_m(t). \qquad S2$$

$\xi_m(t)$ is assumed to have zero mean and a standard deviation of $CV_m\langle L_b\rangle$. $\xi_m(t)$ is assumed to be independent of $L_a(t')$, for all $t'$ within a cell cycle. It is also independent of $\xi_m(t')$ for all $t' \neq t$. The measurement error is estimated using the method previously stated[61]. The method uses the relation between the variance of difference in lengths at time $k\Delta t$ from birth and the length at birth ($Var\big(L(k\Delta t) - L(0)\big)$) vs. time ($k\Delta t$) to obtain the standard deviation of the measurement error. $\Delta t$ is the time interval between consecutive length measurements. We fit the data of $Var\big(L(k\Delta t) - L(0)\big)$ vs. $k\Delta t$ plot to a quadratic equation of the form $y(x) = ax^2 + b$, where $a$ and $b$ are fitting parameters. The parameter $b$ is used to find the measurement error, while $a$ is related to the coefficient of variation of the growth rate. The plots $Var\big(L(k\Delta t) - L(0)\big)$ vs. $k\Delta t$ are shown for the pulse label experiments in acidic pH conditions and unbuffered Mtb data in Extended Data Fig. 6a,b, respectively.

Next, we will show that the measurement errors have a small effect on the binned data trend in the growth rate vs. age and elongation speed vs. age plots. We use simulations where cells grow linearly, and the length at division ($Ld$) is a function of the birth lengths ($Lb$), i.e., $Ld = f(Lb)$. Following Equation $S2$, we also have a size additive measurement noise added to each length measurement collected at equal time intervals in the cell cycle. For details about the simulation, refer to the Simulations section in Methods.

In Extended Data Fig. 6c,d, respectively, we show the binned data trend (in blue) of the growth rate vs. age and elongation speed vs. age plots for a single simulation run of the model. Also shown are the average binned data trend (averaged over 1000 runs) for the plots with the measurement error as calculated in the previous section (solid red line) and without any measurement error (red dashed line). The parameters in the simulations are obtained from the unbuffered Mtb movies.

The Extended Data Fig. 6d clearly shows that the measurement error does not affect the average binned data trend in elongation speed vs. age plot while the effect on growth rate vs. age plots is also negligible (Extended Data Fig. 6c). Thus, these methods are suitable for elucidating the mode of growth.
